# TGFβ signaling is required for tenocyte recruitment and functional neonatal tendon regeneration

**DOI:** 10.1101/767467

**Authors:** Deepak A. Kaji, Kristen L. Howell, Alice H. Huang

**Author notes:** **Corresponding Author:** Alice Huang, PhD, Associate Professor, Dept of Orthopaedics, Icahn School of Medicine at Mount Sinai, 1 Gustave Levy Pl, Box 1188, NY, NY 10029.

## Abstract

Tendon injuries are common with poor healing potential. The paucity of therapies for tendon is due to our limited understanding of the cells and molecules that drive tendon regeneration. Using a model of neonatal mouse tendon regeneration, we determined the molecular basis for regeneration and identify TGFβ signaling as a major pathway. Through targeted gene deletion, small molecule inhibition, and lineage tracing, we elucidated TGFβ-dependent and –independent mechanisms underyling tendon regeneration. Importantly, functional recovery depended on TGFβ signaling and loss of function is due to impaired tenogenic cell recruitment from both *Scx*^*lin*^ and non-*Scx*^*lin*^ sources. We show that TGFβ signaling is required directly in neonatal tenocytes for recruitment and that TGFβ is positively regulated in tendons. Collectively, these results are the first to show a functional role for TGFβ signaling in tendon regeneration and offer new insights toward the divergent cellular activities that may lead to regenerative vs fibrotic healing.

## INTRODUCTION

Tendons connect muscle to bone and function to transmit muscle forces to the skeleton. Tendon function is enabled by a specialized extracellular matrix composed of highly aligned type 1 collagen fibrils (Voleti, Buckley, & Soslowsky, 2012). Although healthy tendon can normally resist high mechanical loads, mechanical properties are permanently impaired after injury due to its minimally regenerative potential (Voleti et al., 2012). This loss of function can lead to chronic pain, decreased quality of life, and increased risk of re-rupture. Current treatment options remain limited and there are almost no cell or biological treatments to improve tendon repair or induce regeneration.

To date, the majority of models for tendon healing are models of scar-mediated healing since adult tendon does not regenerate (Ackerman et al., 2019; Dyment et al., 2014; Dyment et al., 2013; Howell et al., 2017; Katzel et al., 2011; Kim et al., 2011; Mass & Tuel, 1991). Although a few groups showed successful tendon regeneration in model systems such as fetal sheep and MRL/MpJ mice, genetic manipulation is relatively challenging in these model systems (Beredjiklian et al., 2003; Paredes, Shiovitz, & Andarawis-Puri, 2018). To overcome these limitations, we previously established a model of tendon regeneration using neonatal mice, that can be directly compared to adult mice within the same genetic background (Howell et al., 2017). Using lineage tracing, we found that neonatal tendon regeneration is driven by tendon cell proliferation, recruitment, and differentiation leading to full functional restoration. In contrast, adult tendon healing is defined by the persistent presence of myofibroblasts, absence of tendon cell proliferation or recruitment, abnormal cartilage differentiation, and loss of functional properties. Having identified these cellular processes distinguishing neonatal tendon healing from adult, we now focus on the molecular pathways that regulate neonatal tendon regeneration.

Although FGF signaling was first established in chick tendon development (Brent, Schweitzer, & Tabin, 2003), the TGFβ pathway subsequently emerged as the most important signaling pathway identified for mammalian tendon formation (Havis et al., 2016; Kuo, Petersen, & Tuan, 2008; Brian A. Pryce et al., 2009). Signaling through the TGFβ superfamily of molecules is mediated by ligand binding to type II receptors and dimerization with type I receptors. The receptors then phosphorylate intracellular Smad transcription factors that complex with the co-Smad, Smad4, to effect transcriptional change (Shi & Massague, 2003). In mouse embryos, TGFβ ligands are expressed by tendon cells and genetic deletion of the TGFβ type II receptor (TβR2) or TGFβ ligands results in a total loss of tendons (Havis et al., 2016; Kuo et al., 2008; Brian A. Pryce et al., 2009). TGFβ also induces expression of the tendon transcription factor, *Scleraxis* (*Scx*), in a variety of contexts including embryonic limb explants, mesenchymal stem cells, and tendon-derived cells (Brown, Galassi, Stoppato, Schiele, & Kuo, 2015; Maeda et al., 2011; Brian A. Pryce et al., 2009).

In addition to its essential role in tendon development and tendon cell differentiation, TGFβ is also a known inducer of fibrotic scar formation in diverse tissues, including adult tendon (Katzel et al., 2011; Kim et al., 2011; Thomopoulos, Parks, Rifkin, & Derwin, 2015). TGFβ is well established in myofibroblast activation (Border & Noble, 1994; Desmouliere, Geinoz, Gabbiani, & Gabbiani, 1993) and excessive release of TGFβs after injury can also induce tendon cell death (Maeda et al., 2011). Given these contradictory roles of TGFβ signaling in both tendon differentiation and scar formation, it is unclear whether TGFβ signaling enacts a positive or negative response in the context of tendon regeneration. Therefore, in this study, we determined the requirement for TGFβ signaling in neonatal tendon regeneration. Using small molecule inhibition and genetic deletion experiments, we identified a role for TGFβ in functional regeneration and tenogenic cell recruitment from both *Scx* and non-*Scx* lineages.

## RESULTS

### TGFβ signaling is activated after neonatal injury

To determine whether TGFβ signaling is activated after neonatal injury, we evaluated the expression of TGFβ ligands (TGFβ1, 2, 3) as well as the TGFβ type II receptor (TβR2) at d3, d7, d14, and d28 post-injury by qPCR. TβR2 was expressed at all time points after injury and upregulated in injured tendons relative to uninjured controls at d7 and d28 (p < 0.01) (**Figure 1A**). TGFβ1 and TGFβ2 expression levels were transiently upregulated at alternating timepoints, however overall expression levels were quite low for TGFβ2 expression (several fold lower than TGFβs 1 and 3), suggesting a relatively minor role in regeneration for this ligand. By contrast, TGFβ3 expression was increased at all timepoints relative to control (**Figure 1A**). To confirm active TGFβ signaling, western blot was carried out for TβR2 and phospho-Smad2/3. Consistent with gene expression results, we found enhanced TβR2 and phospho-Smad2/3 protein in injured tendons at d14, suggesting active TGFβ signaling (**Figure 1B**). Collectively, these results suggest a potential role for TGFβ signaling in neonatal tendon regeneration and suggest that most of the signaling may be driven by TGFβs 1 and 3.

**Figure 1:**
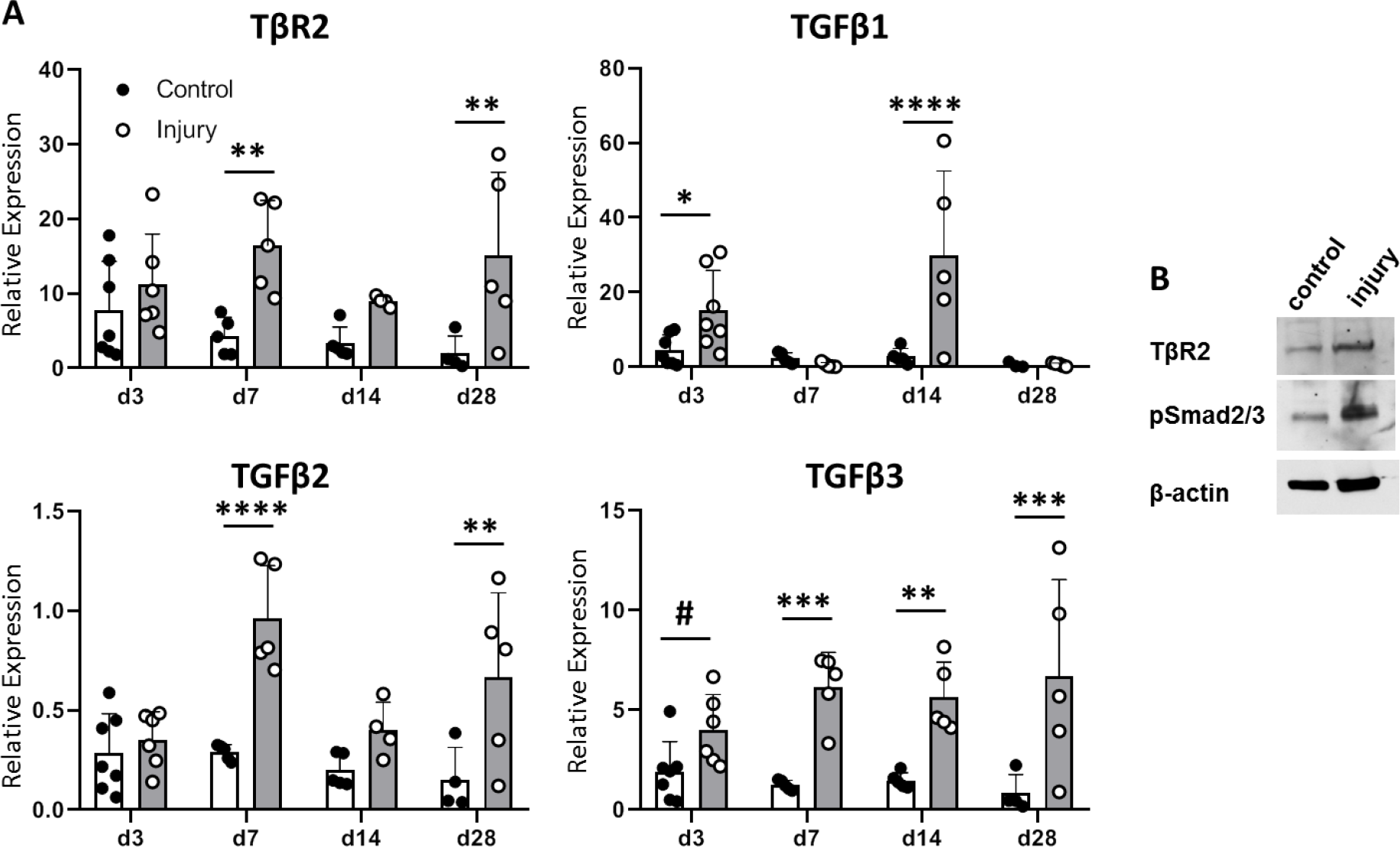
TGFβ signaling is activated after neonatal tendon injury. (**A**) Gene expression of control and injured tendons at d3, d7, d14, and d28 post-injury by qPCR showed upregulation of TGFβ signaling molecules. Expression levels were normalized to *Gapdh* using standard curve method (n=5-7 mice). (**B**) Western blot of control and injured tendons at d14 showed enhanced phospho-Smad2/3 after injury indicating active TGFβ signaling (3 tendons combined). * p<0.05, ** p<0.01, *** p<0.001.

### TGFβ signaling is required for full functional regeneration

To test the requirement for TGFβ signaling in functional restoration, we inhibited TGFβ signaling for 14 days using the small molecule inhibitor SB-431542, which targets the TGFβ family type I receptors ALK 4/5/7. Pups treated with the inhibitor showed no adverse effects in growth compared to carrier-treated pups and tendons appeared grossly normal (**Figure S1**). In a previous study, we determined that the parameters for the brake and propel phases of gait were highly associated with Achilles tendon function (Howell et al., 2017). Carrier-treated mice fully recovered % brake and % propel gait parameters by d14, consistent with functional recovery (**Figure 2A**). By contrast, both % brake and % propel were impaired relative to the contralateral control limb with SB-431542 treatment. We also observed a significant decrease in % propel stride relative to the injured limb of carrier-treated animals. Defects in whole limb gait persisted at d28 for both parameters despite cessation of inhibitor treatment from d14-d28 (**Figure 2B**).

**Figure 2:**
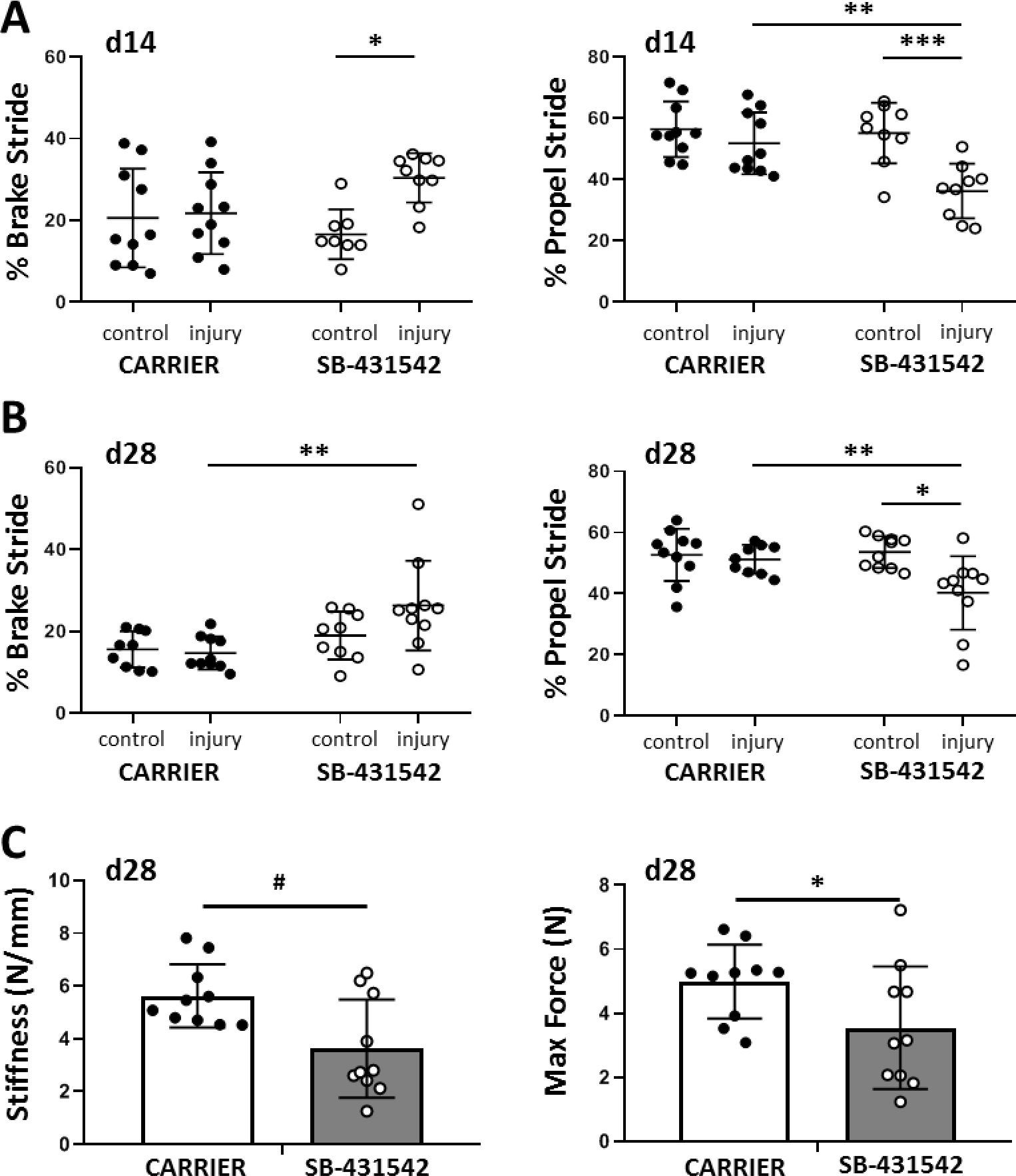
TGFβ signaling is required for functional recovery. Gait analysis at (**A**) d14 and (**B**) d28 showed impaired % brake stride and % propel stride after injury with SB-431542 treatment. (**C**) Tensile testing revealed reduced stiffness and max force with SB-431542 treatment. # p<0.1, * p<0.05, ** p<0.01, *** p<0.001 (n=8-10 mice).

To determine the mechanical properties of the healing tissue directly, we then performed tensile testing of the tendons at d28 and observed a reduction in stiffness and max force with SB-431542 treatment (**Figure 2C**). Mechanical properties in uninjured control tendons were not significantly different, further indicating that postnatal growth was not impaired with TGFβ inhibition (**Figure S1**). Taken together, these data show that TGFβ is required in the first 14 days of healing for functional regeneration.

### TGFβ signaling in neonatal tenocytes is required for cell recruitment after injury

We previously found that *Scx*-lineage (*Scx*^*lin*^) tenocyte proliferation, recruitment, and differentiation are unique features of the neonatal regenerative response that are not observed during adult healing (Howell+, Sci Rep, 2017). To first assess whether tenocyte recruitment is affected by TGFβ inhibition, we used the *ScxCreERT2* mouse to genetically label differentiated tenocytes prior to injury and traced the fate of these cells during healing when TGFβ signaling is suppressed. In carrier-treated mice, whole mount imaging of hindlimbs showed *Scx*^*lin*^ cells (*RosaT*+) occupying the gap space between the original Achilles tendon stubs at d14 (**Figure 3A**), while little *RosaT* signal was detected in SB-431542-treated limbs (**Figure 3B**). Quantification of transverse sections taken from the midsubstance regions confirmed reduced *Scx*^*lin*^ tenocyte numbers with TGFβ inhibition (**Figure 3C, 3D**).

**Figure 3:**
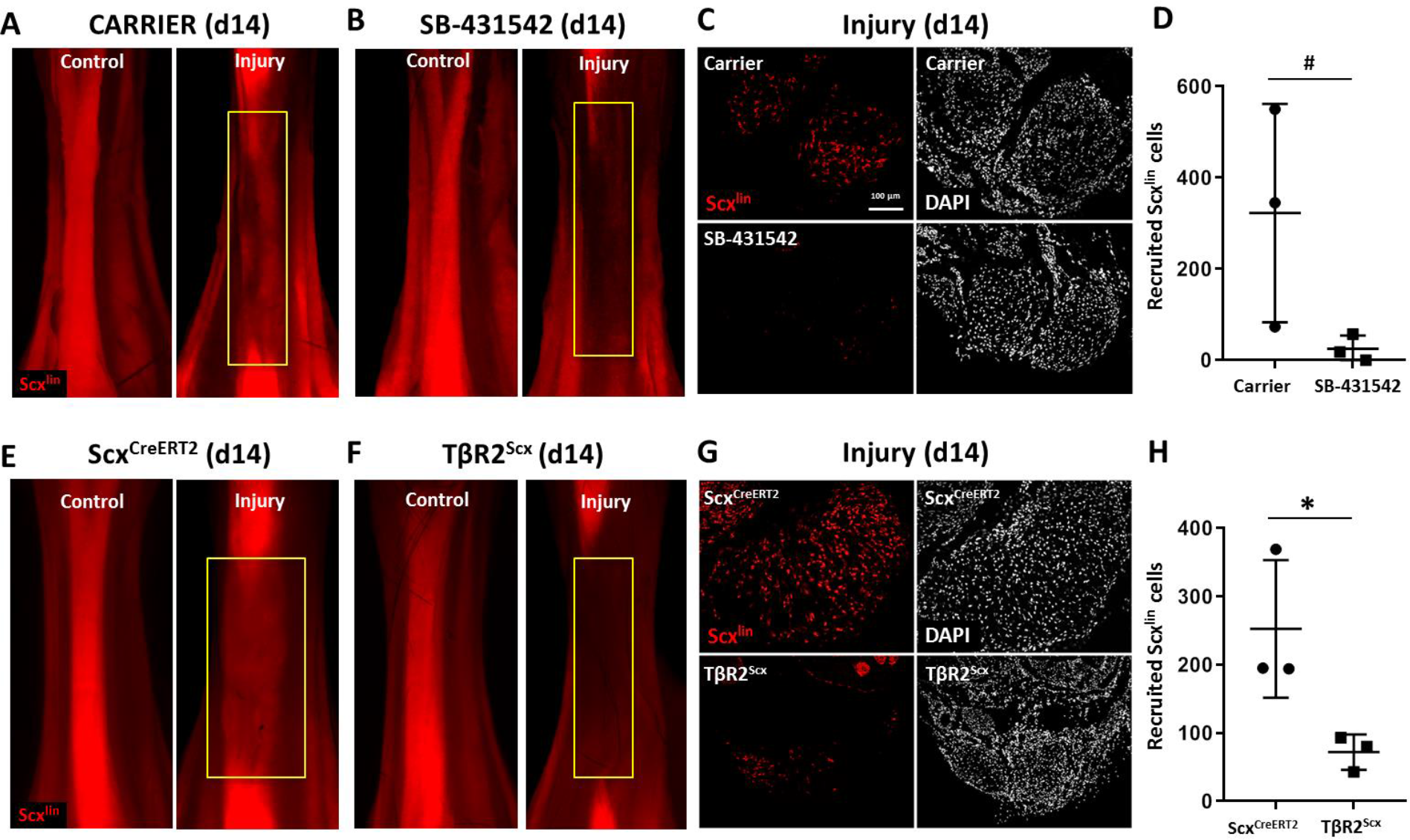
TGFβ signaling is required for tenocyte recruitment. (**A**, **B**) Whole mount images of control and injured limbs in carrier-treated and SB-431542-treated *Scx*^*CreERT2*^/*RosaT* mice. (**C**) Transverse sections through the neo-tendon (yellow boxes) near the mid-substance and (**D**) quantification showed reduced *Scx*^*lin*^, *RosaT*+ cell recruitment with SB-431542 treatment (n=3 mice). (**E**, **F**) Whole mount images of control and injured limbs in wild type *Scx*^*CreERT2*^/*RosaT* and *TβR2*^*Scx*^/*RosaT* mice. (**G**) Transverse sections through the neo-tendon near the mid-substance and (**H**) quantification showed reduced *Scx*^*lin*^, *RosaT*+ cell recruitment in *TβR2*^*Scx*^ mutants (n=3 mice). * p<0.05, # p<0.10. Scalebars: 100 μm.

Since SB-431542 treatment indiscriminantly targets all cells, we next tested whether neonatal tenocytes directly required TGFβ signaling for their recruitment. We therefore deleted *TβR2* using *ScxCreERT2* prior to injury (*TβR2*^*Scx*^) and visualized mutant cells by *RosaT* expression. Since TGFβ signaling is mediated by a single type II receptor (*TβR2*), all TGFβ signaling is abolished with deletion of this receptor. Consistent with our inhibitor studies, few *Scx*^*lin*^ tenocytes were detected in the midsubstance of *TβR2*^*Scx*^ mutant tendons compared to *ScxCreERT2* wild type tendons (**Figure 3F-3H**). This data suggests that TGFβ signaling is required in neonatal tenocytes for recruitment, rather than an indirect effect of TGFβ inhibition.

### TGFβ signaling is required for tenocyte migration but not proliferation

We hypothesized that the absence of *Scx*^*lin*^ cell recruitment at d14 with TGFβ inhibition may be due to a defect in cell proliferation at an earlier timepoint. In a previous study, we showed intense *Scx*^*lin*^ tenocyte proliferation that was localized at the cut site of tendon stubs at d3. To test this hypothesis, we collected *ScxCreERT2*-labeled limbs at d3 post-injury with SB-431542 treatment as well as *TβR2*^*Scx*^ deletion. Consistent with previous findings, transverse sections through the midsubstance gap space confirmed that *Scx*^*lin*^ cells were not detectable at d3 after injury for any condition (not shown). EdU staining of proliferating *Scx*^*lin*^ tendon cells showed comparable numbers between carrier-treated and SB-431542-treated mice after injury (**Figure 4A-C**). Similarly, no differences were detected between injured, wild type and *TβR2*^*Scx*^ mutants (**Figure 4D-F**). At this timepoint, tenocyte proliferation in uninjured control Achilles tendons was extremely low (0-1 EdU+/*Scx*^*lin*^+ cell per section) and was unaffected by SB-431542 treatment or *TβR2*^*Scx*^ deletion (**Figure S2**).

**Figure 4:**
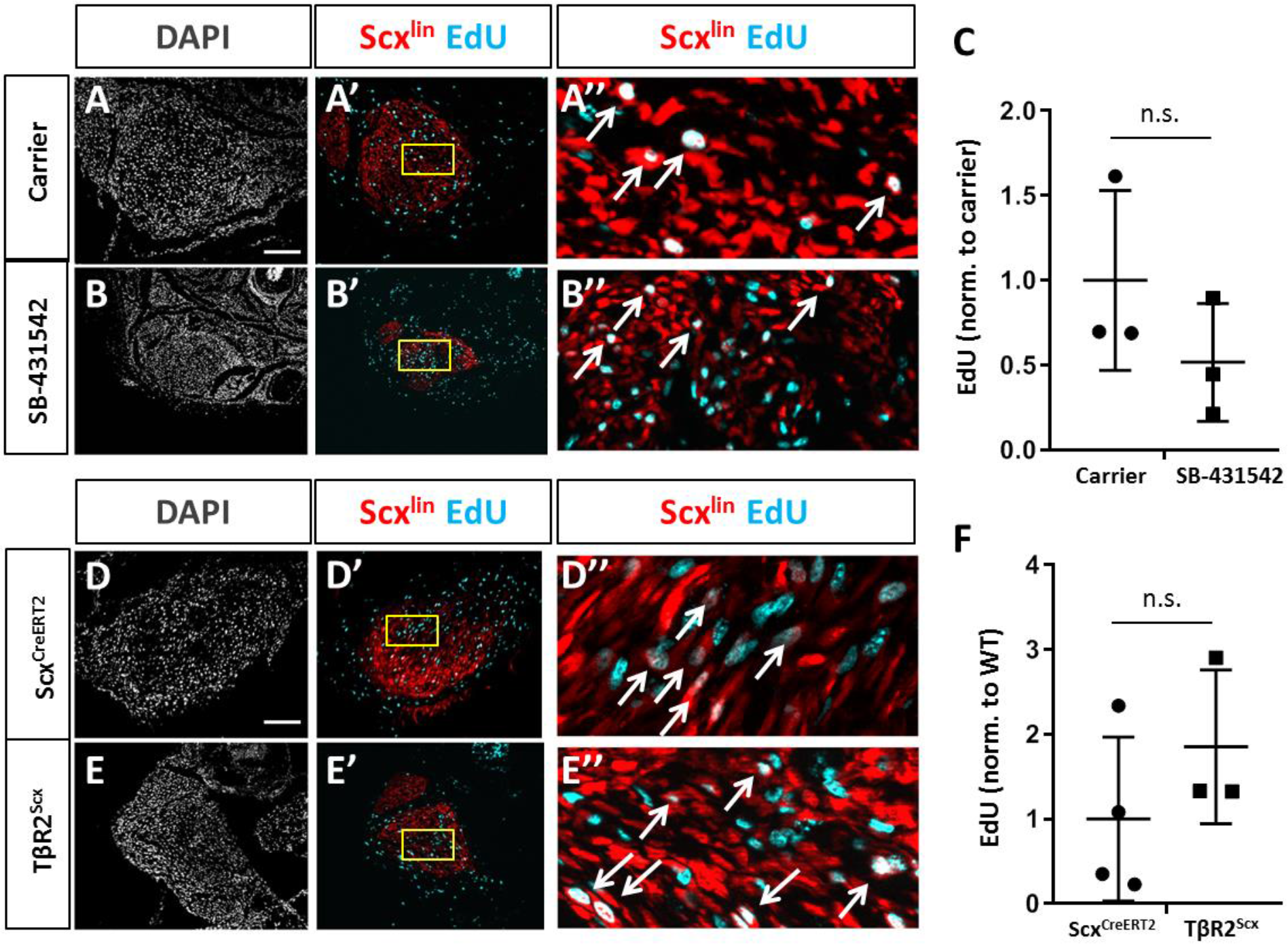
TGFβ signaling is not required for tenocyte proliferation. Transverse sections through the cut site of (**A**, **A′**, **A″**) carrier-treated injured tendon or (**B**, **B′**, **B″**) SB-431542-treated injured tendon stained for EdU and counterstained with DAPI. A″, B″, D″, E″ are enlarged images from yellow boxed regions shown in A′, B′, C′, D′. (**C**) Quantification of EdU and *Scx*^*lin*^ overlays showed no difference in *Scx*^*lin*^ cell proliferation after injury with TGFβ inhibition(n=3 mice). Transverse sections through the cut site of (**D**, **D′**, **D″**) wild type injured tendon or (**E**, **E′**, **E″**) *TβR2*^*Scx*^ injured tendon stained for EdU and counterstained with DAPI. (**F**) Quantification of EdU and *Scx*^*lin*^ overlays show no difference in *Scx*^*lin*^ cell proliferation after injury with *TβR2* deletion (n=3 mice). White arrows indicate EdU+, *Scx*^*lin*^ cells. n.s. indicates p>0.1. Scalebars: 100 μm.

Since proliferation was not affected, we next determined whether TGFβ signaling may be required for tenocyte migration. *In vitro* wounds were created in cell monolayers and migration of cells into the defect observed over 12 hours (**Figure 5A**). Tenocytes in DMEM alone did not migrate at any timepoint. Differences in wound closure were not observed between DMEM and DMEM+FBS until 12 hours. In contrast, the addition of TGFβ1 significantly enhanced cell migration, and differences in wound closure were detected as early as 4 hours with nearly full wound closure by 8 hours (**Figure 5A, B**).

**Figure 5:**
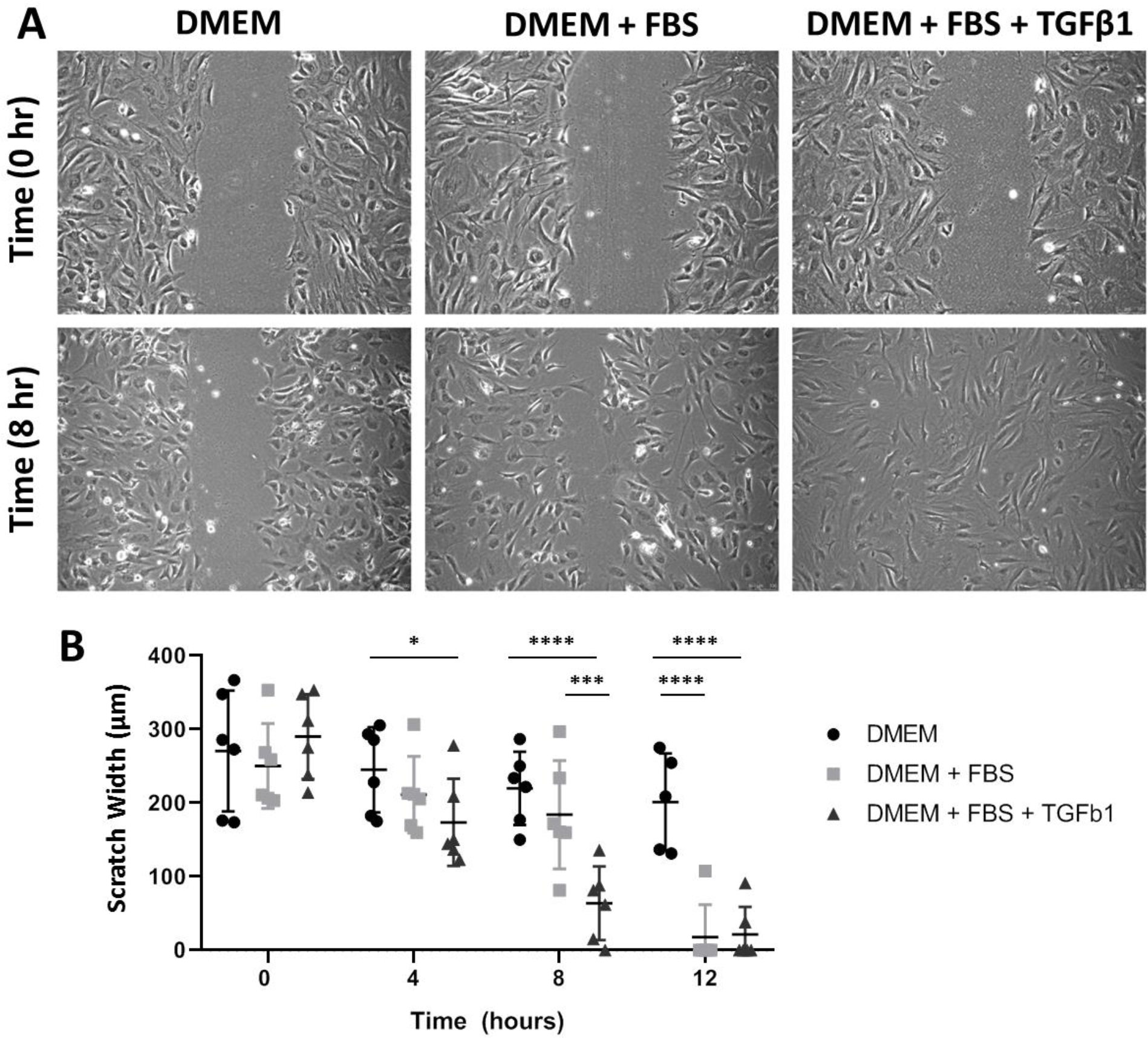
TGFβ enhances neonatal tenocyte migration *in vitro*. (*A*) Phase contrast images and (*B*) quantification of *in vitro* wound assay show rapid closure with TGFβ1 supplementation relative to DMEM and DMEM+FBS conditions (n=6). * p<0.05, *** p<0.001, **** p<0.0001.

Collectively, these results suggest that TGFβ signaling is required in neonatal tenocytes for cell recruitment, and that recruitment occurs by active cell migration rather than growth through proliferation.

### Increased TGFβ ligand production in injured tendon depends on TGFβ signaling

Although *Scx*^*lin*^ cells are not present in the gap space at d3, the region is not devoid of cells. At this time, we observed early accumulation of αSMA+ cells that are not from the *Scx*^*lin*^ (Howell et al., 2017). Surprisingly, immunostaining for αSMA revealed that recruitment of αSMA+ cells at d3 was not affected by TGFβ inhibition or *TβR2* deletion (**Figure 6A, B**). Transverse sections through the midsubstance gap space also confirmed that *Scx*^*lin*^ cells were not yet detectable at d3 in any condition (not shown). The presence of αSMA+ cells within the gap space prior to *Scx*^*lin*^ cell recruitment suggested that these cells may be a source of TGFβ ligands that signal to tenocytes for migration. Immunostaining for all three TGFβ isoforms showed comparable levels of signal between gap space cells and control tendon (**Figure S3**). We considered the possibility that ligand production may be regulated by TGFβ signaling and that reduced presence of TGFβ ligands may prevent tenocyte recruitment. However, immunostaining for TGFβ ligands showed equivalent staining intensity within the gap space for carrier- and SB-431542-treated limbs (**Figure S3**). Differences were only observed in injured tendons; we found increased staining of ligands for carrier-treated injured tendons, but this increase was not detected with SB-431542 treatment (**Figure 7A, B**). This data suggests that TGFβ signaling is required for upregulation of TGFβ ligands with injury, and that this is likely autonomously regulated in tenocytes.

**Figure 6:**
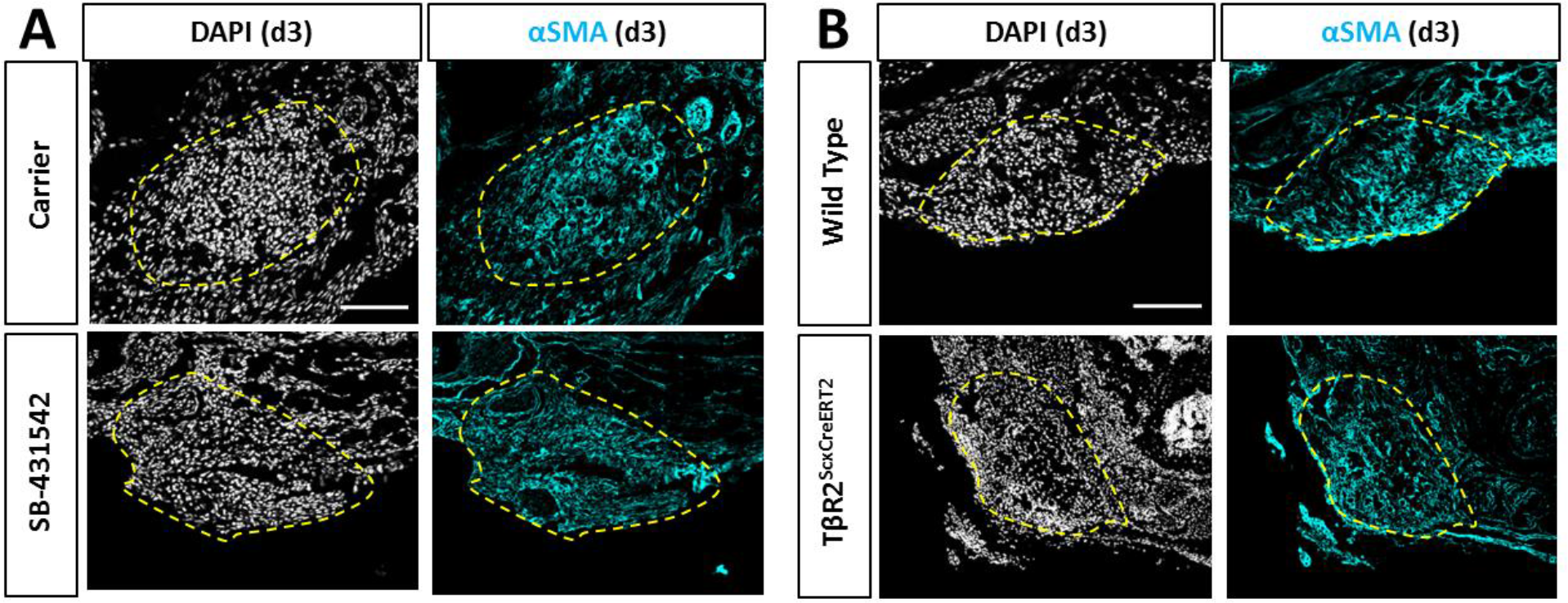
Recruitment of αSMA+ myofibroblasts is not affected by TGFβ inhibition or *TβR2*^*Scx*^ deletion. Transverse sections through the gap space at d3 showed abundant αSMA+ cells with (**A**) SB-431542 treatment or (**B**) *TβR2*^*Scx*^ deletion at levels comparable to carrier-treated or wild type. Yellow dashed outlines highlight gap area formerly occupied by the Achilles tendon.

**Figure 7:**
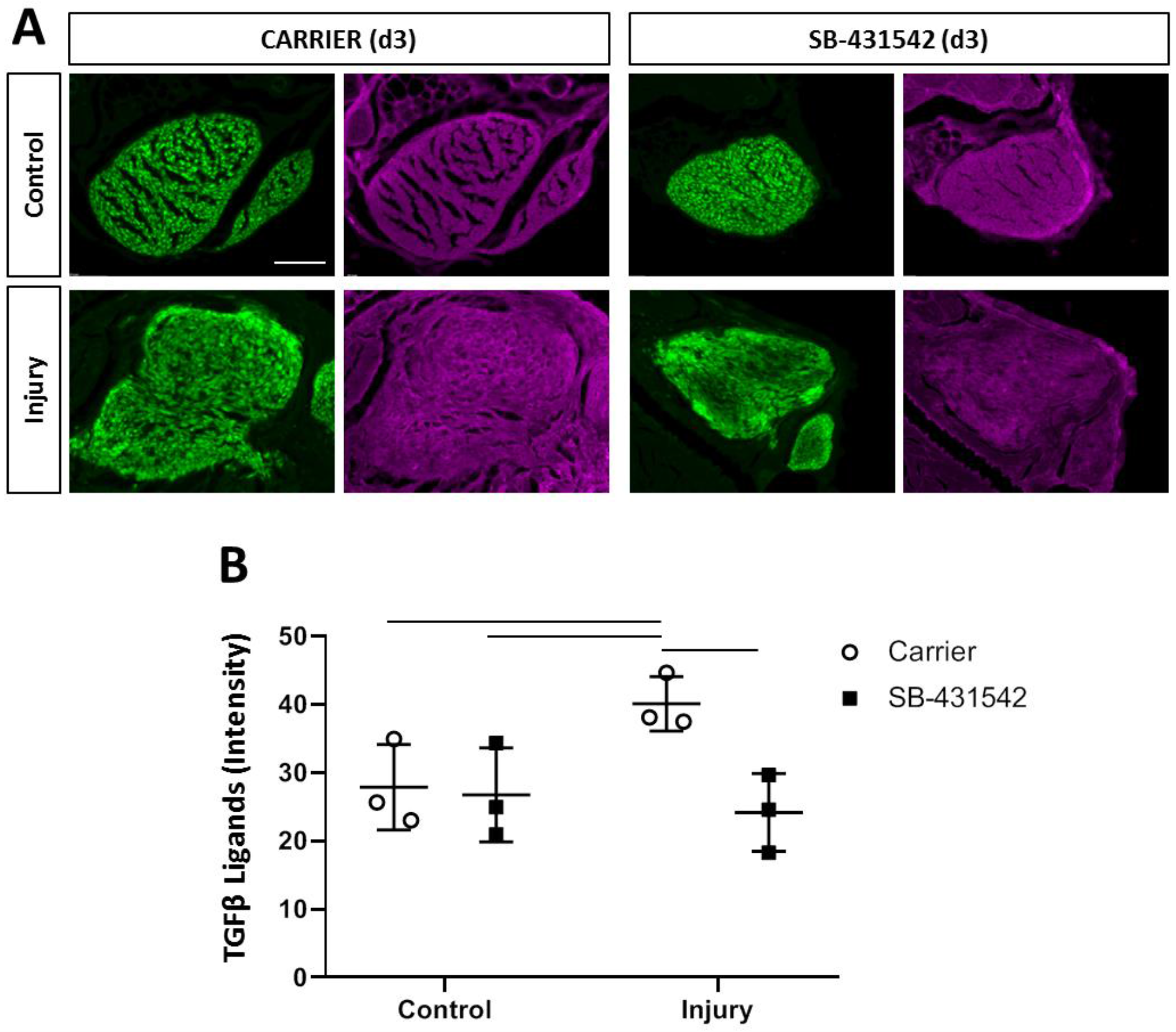
TGFβ ligand synthesis after injury is regulated by TGFβ signaling. (**A**) Transverse sections through the tendon at d3 immunostained with antibody against all three TGFβ isoforms. (**B**) Quantification of intensity levels show increased TGFβ ligands after injury in carrier-treated tendons that is no longer observed with SB-431542 treatment. Bars indicate p<0.05. Scalebar: 100 μm.

### Non-*Scx*^*lin*^ tenogenic cells also contribute to neotendon formation

Although αSMA+ cells are present at d3, immunostaining showed few αSMA+ cells by d14 for all experimental conditions (**Figure S4**), consistent with our previous study (Howell et al., 2017). We hypothesized that αSMA+ cells (which are not *Scx*^*lin*^) may differentiate toward the tendon lineage and turn off αSMA. This is supported by previous studies using *αSMACreERT2*, which showed that *αSMA*^*lin*^ cells of the paratenon turn on *ScxGFP* with adult patellar tendon injury (Dyment et al., 2013). Analysis of *ScxGFP* expression in carrier-treated injured limbs indeed showed a population of non-*Scx*^*lin*^, *ScxGFP*+ cells comprising the neo-tendon (**Figure 8A, B**). Comparison to contralateral non-injured controls indicated that incomplete recombination of *Scx*^*lin*^ cells does not explain this phenomenon since recombination efficiency is ~96.4% in control tendons. Quantification of the non-*Scx*^*lin*^ *ScxGFP*+ (*ScxGFP* only) population showed fewer *ScxGFP* only cells in the neo-tendon after injury in SB-431542-treated mice (**Figure 8C-E**). There was a proportional decrease in DAPI+ cells, indicating that the reduction in *ScxGFP* only cells was probably not due to failure of cells within the gap space to differentiate. Rather, the cells that comprise this population are either not recruited or experience reduced proliferation in the absence of TGFβ signaling. To determine whether these non- *Scx*^*lin*^, *ScxGFP*+ cells were derived from αSMA+ cells, we used the transgenic *αSMACreERT2* mouse and labeled cells by tamoxifen administered at P2, P3. Analysis of transverse cryosections at P5 showed an unexpected amount of recombination in *ScxGFP* tenocytes (**Figure S5**). Immunostaining with anti-αSMA confirmed that neonatal tenocytes normally do not express αSMA. The surprising extent of tendon cell recombination with the *αSMACreERT2* therefore precluded its use in identifying the source of non-*Scx*^*lin*^ *ScxGFP*+ cells after injury.

**Figure 8:**
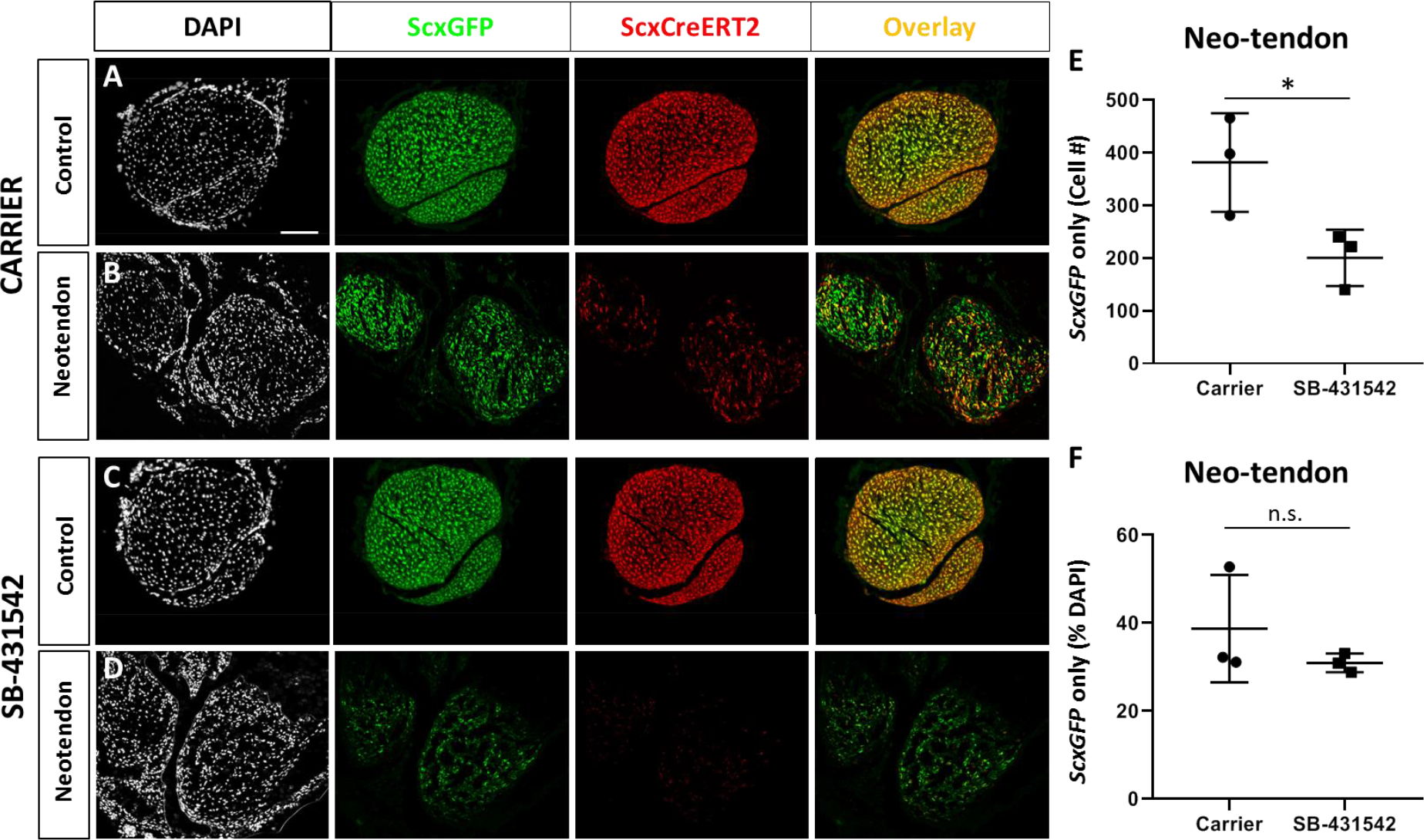
TGFβ ligand synthesis after injury is regulated by TGFβ signaling. Transverse sections through the neo-tendon of control and injured limbs in (**A, B**) carrier-treated and (**C, D**) SB-431542-treated *Scx*^*CreERT2*^/*RosaT*/*ScxGFP* mice. (**E, F**) Quantification of non-*Scx*^*lin*^, *ScxGFP*+ cells show reduction in cell number with SB-431542 treatment but not when normalized to total DAPI+ cells (n=3 mice). * p<0.05. n.s. indicates p>0.1. Scalebar: 100 μm.

To test whether tendon-specific differentiation is affected, we also determined gene expression by real time qPCR at d3 and d14 post-injury. At d3, injured limbs from carrier-treated mice decreased tendon markers *Scx*, *Mkx* and *Tnmd* compared to their contralateral uninjured controls (**Figure 9A**).

**Figure 9:**
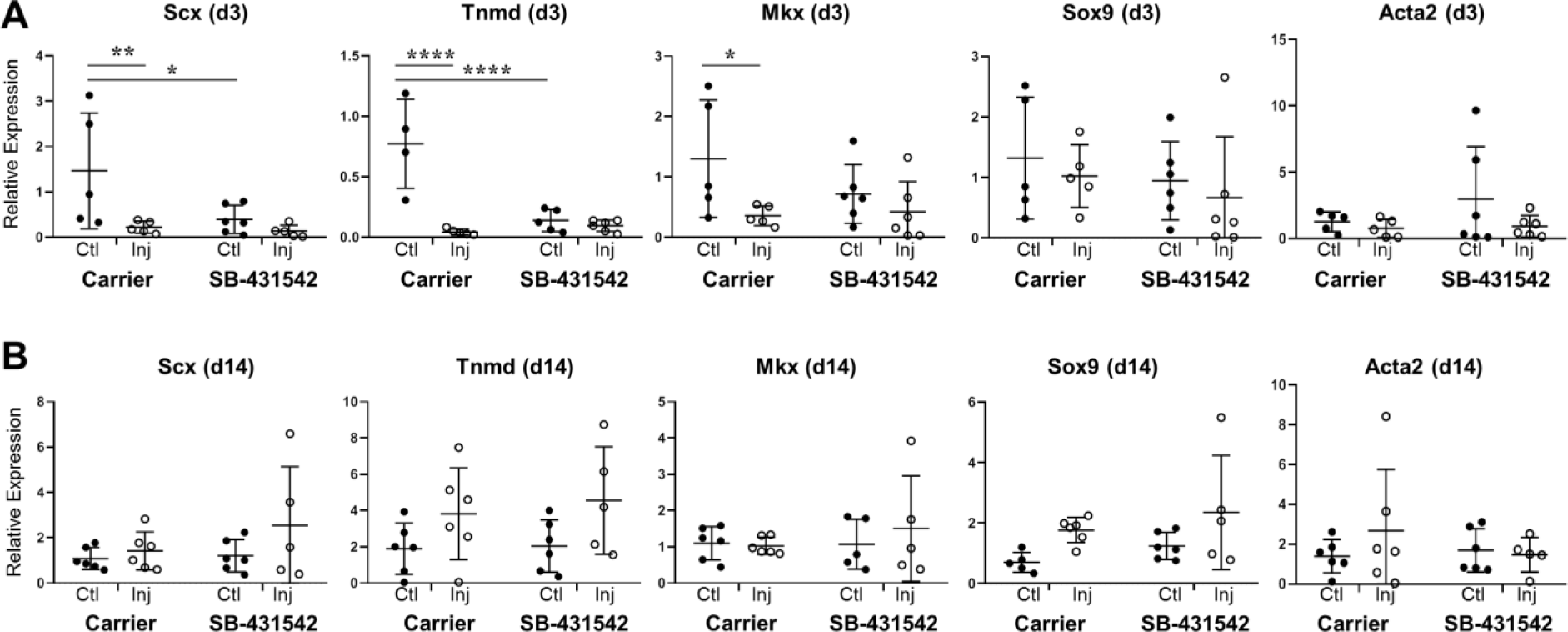
Tendon gene expression is not affected by TGFβ inhibition at d14 post-injury. Real time qPCR analysis of tendons harvested at (**A**) d3 and (**B**) d14 from carrier-treated and SB-431542-treated animals (n=4-6 mice). Tendon genes were decreased after injury in carrier-treated mice but not with SB-431542 treatment at d3. Differences were no longer detected by d14. Cartilage and myofibroblast markers were not significantly different. * p < 0.05, ** p<0.01, **** p<0.0001.

Interestingly, SB-431542-treatment also decreased tendon gene expression in uninjured control tendons relative to carrier controls. Additional decreases with injury were not detected between injured and control tendons with TGFβ inhibition. By d14, tendon gene expression was similar across all samples regardless of treatment or injury (**Figure 9B**). Expression levels of *Sox9* and *Acta2* (the gene for αSMA) were not different across experimental conditions at either timepoint (**Figure 9A, B**). These data indicate that despite defects in tenogenic cell recruitment, tendon gene expression after injury was largely not affected by TGFβ inhibition.

Taken together, our results reveal TGFβ-dependent and TGFβ-independent processes during neonatal tendon regeneration. While early proliferation of *Scx*^*lin*^ tenocytes and activation of extrinsic aSMA+ myofibroblasts do not depend on TGFβ, subsequent recruitment of tenogenic cells (from *Scx*^*lin*^ and non-*Scx*^*lin*^ sources) and functional restoration depend on TGFβ signaling (**Figure 10**).

**Figure 10:**
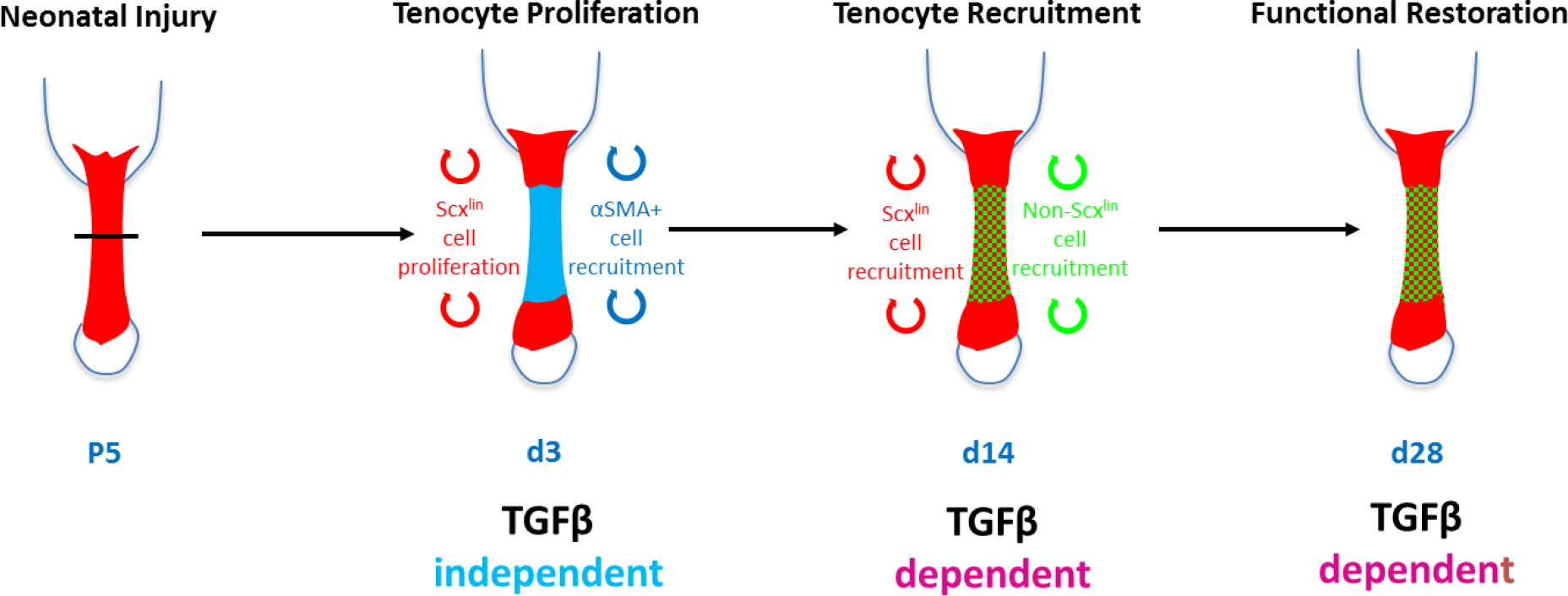
Requirement for TGFβ signaling in neonatal regeneration. Conceptual model schematic depicting the TGFβ-dependent and TGFβ-independent cellular processes during neonatal tendon regeneration. While tenocyte proliferation and αSMA cell recruitment at d3 occur independently of TGFβ signaling, tenogenic cell recruitment and functional restoration at subsequent timepoints require TGFβ.

## DISCUSSION

TGFβ signaling is a known regulator of many cellular processes, including proliferation, survival, migration, and differentiation (Shi & Massague, 2003). In tendon, TGFβ signaling is essential in embryonic tendon development as well as induction and maintenance of tendon cell fate (Brian A. Pryce et al., 2009). However, this pathway is also strongly identified with fibrotic, scar-mediated healing and excessive TGFβ signaling results in tenocyte apoptosis (Davies et al., 2016; Katzel et al., 2011). In the context of tendon regeneration, it was therefore unclear whether TGFβ would be required for tenogenesis or whether activation of TGFβ would drive fibrotic responses. Using our previously established model of neonatal tendon regeneration, we now show that TGFβ signaling is enhanced after injury and is required in neonatal tenocytes for their recruitment. Since tenocyte proliferation was not affected, we propose that tenocyte-mediated regeneration requires active migration of cells to bridge the gap space. This is further supported by *in vitro* data showing enhanced migrational capacity of neonatal tenocytes in the presence of TGFβ ligand and is consistent with several studies in the literature for other cell types (Shi & Massague, 2003).

In addition to intrinsic tenocytes, we also identified a second population of non-*Scx*^*lin*^, *ScxGFP*+ cells that are also recruited to the gap space. Inhibition of TGFβ signaling also resulted in reduced numbers of these cells. One potential source of these cells may be the epitenon as it was previously proposed that tendon stem/progenitor cells reside in epitenon (Dyment et al., 2014; Dyment et al., 2013; Gumucio, Phan, Ruehlmann, Noah, & Mendias, 2014; Mendias, Gumucio, Bakhurin, Lynch, & Brooks, 2012; Mienaltowski, Adams, & Birk, 2013). Although lineage tracing with *αSMACreERT2* showed restricted labeling in the epitenon/paratenon in adults (Dyment et al., 2014), we found considerable labeling in tenocytes at neonatal stages which precluded the use of this line to target epitenon-derived cells. Another source may be nearby vasculature, as CD146+ pericytes have been identified for tendon (Lee et al., 2015). Despite impaired recruitment of tenogenic cells with TGFβ inhibition, the expression of tenogenic markers *Scx*, *Tnmd*, and *Mkx* were not different at d14. Identifying additional markers for tendon cell fate is the focus of ongoing studies.

Unexpectedly, we found that early activation of αSMA+ myofibroblasts is not affected when TGFβ signaling is inhibited. While myofibroblast phenotypes were not expected in *TβR2*^*Scx*^ mutants (since *Scx* lineage tracing showed that myofibroblasts did not derive from tenocytes), we were surprised to observe abundant myofibroblast accumulation with global SB-431542 inhibition. Since TGFβ signaling is well-established in myofibroblast induction and survival, these results suggest that our inhibition protocol likely did not abrogate all TGFβ signaling and there may be different threshholds required for TGFβ-dependent myofibroblast activation versus tenocyte recruitment. Alternatively, other signaling pathways have also been implicated in myofibroblast activation in the absence of TGFβ, including CTGF, EGF, and IGF2 (Grotendorst, Rahmanie, & Duncan, 2004). Interestingly, CTGF can induce tendon differentiation of adipose derived stem cells and delivery of CTGF improves adult tendon healing by activating endogenous stem cells (Lee et al., 2015; Thomopoulos et al., 2015). Identifying the role of CTGF and other pathways in neonatal regeneration may provide additional insights in poor adult tendon healing.

We identified a potential source of TGFβ ligands in myofibroblasts within the gap space, which may drive directional migration of the tenocytes from the stubs. Although abundant TGFβs were also detected in the tendon matrix of uninjured controls, these ligands may be in an inactive state since TGFβs are typically secreted in a latent form bound to the extracellular matrix. Release of TGFβs to its active form can be induced by proteases or mechanically (such as with transection injury) (Maeda et al., 2011). We detected an increase in tendon TGFβ ligands after injury, which was suppressed by small molecule inhibition of TGFβ signaling. This suggests that initiation of TGFβ signaling (possibly by release of TGFβs from the matrix with transection) results in positive feedback in tenocytes. Other sources of TGFβs may be immune cells, which are also known to produce TGFβs. Of the three TGFβ isoforms, gene expression data suggested that the primary ligands driving neonatal regeneration may be TGFβs 1 and 3. Although TGFβ1 showed bimodal upregulation pattern, TGFβ3 was consistently upregulated after injury. During embryonic development, TGFβs 2 and 3 are expressed in tendons and allelic deletion of these ligands results in increasing loss of tendons (Kuo et al., 2008; Brian A. Pryce et al., 2009); in the context of injury, TGFβ3 is expressed during regenerative fetal tendon healing in sheep while TGFβ1 is associated with fibrotic adult tendon healing (Beredjiklian et al., 2003; Kim et al., 2011). Although this supports the notion that TGFβs 2 and 3 are pro-tenogenic relative to TGFβ1, it is unclear whether the individual ligands actually can activate distinct healing or tenogenic responses. Additional research must therefore be carried out to elucidate their activities.

Although adult tendon healing was not determined in this study, it is well established that TGFβ signaling is elevated after adult injury and results in fibrotic scar formation. Inhibition of TGFβ signaling, either with neutralizing antibodies or via *Smad3*^−/−^ deletion attenuates fibrosis but fail to regenerate tendon structure or function (Katzel et al., 2011; Kim et al., 2011). We previously showed that adult tenocytes are largely quiescent after full transection injury with minimal cell proliferation or recruitment. The distinctive response of neonatal vs adult tenocytes to TGFβ may reflect differences in intrinsic potential (for example adult tenocytes are post-mitotic) or the activation of other signaling pathways that may interact with or modify TGFβ signaling. In addition to Smad signaling, TGFβs can also activate a number of non-Smad pathways; there may be differences in downstream signaling between neonatal and adult tenocytes. Using an *in vitro* engineered tendon model, we previously showed that the tenogenic outcomes of TGFβ signaling did not depend on *Smad4* (Chien, Pryce, Tufa, Keene, & Huang, 2018). Whether this finding is applicable in the context of *in vivo* injury remains to be determined.

Interestingly, while adult tenocytes fail to undergo tenogenic recruitment, a subset of adult tenocytes differentiate along the cartilage lineage, followed by heterotopic ossification (HO) (Howell et al., 2017). This process is not observed during neonatal tendon healing. Inhibition of BMP signaling reduces HO formation in adult tendons, however HO is not completely abolished (Zhang et al., 2016). Although TGFβ is a strong tendon inducer, it is also widely used to induce chondrogenesis in mesenchymal stem cells. During embryonic development, TGFβ also induces a bipotent population of *Scx*+/*Sox9*+ progenitor cells that subsequently contribute to the cartilage and tendon cells of the tendon-skeletal attachment (Blitz, Sharir, Akiyama, & Zelzer, 2013). Whether TGFβ signaling may also play a role in adult tendon HO will be the focus of future studies.

## METHODS

### Experimental procedures

The following mouse lines were used: *ScxGFP* tendon reporter (B. A. Pryce, Brent, Murchison, Tabin, & Schweitzer, 2007), *ScxCreERT2* (generated by R. Schweitzer), *αSMACreERT2* (Grcevic et al., 2012), *Ai14 Rosa26-TdTomato* Cre reporter (Madisen et al., 2010), and *TβR2*^*f*/*f*^ (Chytil, Magnuson, Wright, & Moses, 2002). Lineage tracing and Cre deletion was performed by delivering tamoxifen in corn oil to neonatal mice at P2 and P3 (1.25 mg/pup) (Howell et al., 2017). EdU was given at 0.05 mg 2 hours prior to harvest to label proliferating cells. Global TGFβ inhibition was carried out using the well-established small molecule inhibitor SB-431542 (10 mg/kg, intraperitoneal injection) which targets the TGFβ family type I receptors ALK 4/5/7 (Hamilton, Foster, & Bonnet, 2014; Inman et al., 2002; Laping et al., 2002; Lemos et al., 2015; Waghabi et al., 2009). Daily injections of SB-431542 were administered from day 0-14 after injury. Full Achilles tendon transection without repair was carried out in neonates at P5, with male and female mice distributed evenly between groups. All procedures were approved by the Institutional Animal Care and Use Committee at Mount Sinai.

### Migration assay

Neonatal tenocytes were isolated from P7 pups by digestion in 1% collagenase type 1 (Cat. # LS004188, Worthington, Lakewood, NJ) and collagenase type 4 (Cat. # LS004188, Worthington, Lakewood, NJ) for 4 hours. Cells were expanded and maintained in high glucose DMEM (Cat. # 11965092, Life Technologies, Carlsbad, CA) with 10% fetal bovine serum (FBS, Life Technologies, Carlsbad, CA) and 1% PenStrep (Life Technologies, Carlsbad, CA). For the migration assay, cells were maintained in DMEM only, DMEM+10% FBS, or DMEM+10% FBS+10 ng/mL TGFβ1 (Cat. # 240-B, R&D Systems, Minneapolis, MN). A P200 tip was used scratch down the midline of every well. Phase contrast images were then taken every 4 hours for a total of 12 hours.

### Whole mount fluorescence imaging

Hindlimbs were fixed in 4% paraformaldehyde (PFA, Cat. # 50-980-495, Fisher Scientific, Waltham, MA) overnight at 4°C and skin removed to expose the Achilles tendon. Whole mount images of the posterior limbs were captured using a Leica M165FC stereomicroscope with fluorescence capabilities. Exposure settings were maintained across limbs.

### Immunofluorescence and microscopy

After sacrifice, limbs were fixed in 4% PFA for 24 hours at 4°C, decalcified in 50 mM EDTA for 1-2 weeks at 4°C, then incubated in 5% sucrose (1 hour) and 30% sucrose (overnight) at 4°C. Limbs were then embedded in optimal cutting temperature medium (Cat. # 23-730, Fisher Scientific, Waltham, MA) and 12 um transverse cryosections obtained. Immunostaining was carried out with primary antibodies against αSMA (Cat. # A5228, Sigma, St. Louis, MI), TGFβ 1,2,3 ligands (Cat. # MAB1835, R&D Systems, Minneapolis, MN) and secondary detection with antibodies conjugated to Cy5 (Cat. # 711-175-152; 016-170-084, Jackson ImmunoResearch, West Grove, PA). EdU labeling was detected with the Click it EdU kit in accordance with manufacturer’s instructions (Cat. # C10340, Life Technologies, Carlsbad, CA). Fluorescence images were acquired using the Zeiss Axio Imager with optical sectioning by Apotome or using the Leica DMB6 microscope. Cell quantification was performed in Zeiss Zen or Image J software on transverse cryosection images. All images for quantifications were taken at the same exposure and image manipulations applied equally across samples.

### RNA isolation, reverse transcription, and qRT-PCR

Trizol/chloroform extraction was used to isolate RNA from dissected tendons. cDNA was then synthesized via reverse transcription using the SuperScript VILO master mix (Cat. # 11755050, Invitrogen, Carlsbad, CA). Gene expression was assessed by qRT-PCR using SYBR PCR Master Mix (Cat. # 4309155, Thermo Fisher, Waltham, MA) and calculated using the standard curve method or the 2^−∆∆Ct^ method relative to carrier-treated control tendons. The housekeeping gene, *Gapdh*, was used to normalize gene expression. Primer sequences for TGFβ-related molecules are listed in **Supplemental Table 1**. All other primers were previously described (Howell+, Sci Rep, 2017).

### Gait analysis

Mice were gaited at 10 cm/s for 3-4 s using the DigiGait Imaging System (Mouse Specifics Inc., Quincy, MA). A high-speed digital camera was used to capture forelimb paw positions and parameters previously established for mouse Achilles tendon injury were then extracted (Howell+, Sci Rep, 2017). All parameters were normalized to Stride length to account for differences in animal size and age.

### Biomechanical testing

Tensile testing was performed in PBS at room temperature using custom 3D printed grips to secure the calcaneus bone and Achilles tendon (Abraham et al., 2019). Tendons were preloaded to 0.05N for ~1 min followed by ramp to failure at 1%/s. Structural properties were recorded; since cross-sectional area could not be accurately measured due to the small size of the tissues, material properties were not analyzed.

### Statistical analysis

Quantitative results are presented as mean±standard deviation. Two way ANOVA was used for comparisons with two independent variables (injury and TGFβ inhibition); where significance was detected, posthoc testing was then carried out (Graphpad Prism). All other quantitative analyses were analyzed using Students t-tests. Significant outliers were detected using Grubb’s test (Graphpad Prism). Sample sizes for gait and mechanical properties quantification were calculated from power analyses with power 0.8 and 5% type I error. Samples sizes for other quantitative data were used based on previous data from the lab and published literature. Significance was determined at p<0.05.

## ACKNOWLEDGMENTS

This work was supported by NIH/NIAMS R01AR069537 to AHH and F31AR073626 to DK. We also would like to acknowledge Bhavita Walia for experimental assistance.

## COMPETING INTERESTS

There are no competing interests

## SUPPLEMENTAL DATA

**Supplemental Table 1:**
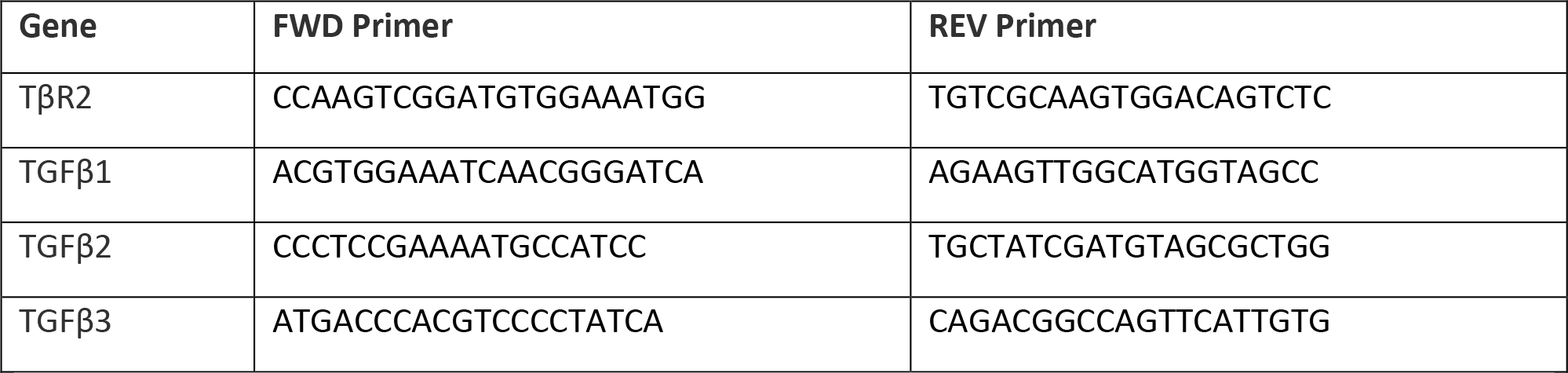
Primer sequences for real time qPCR.

**Figure S1:**
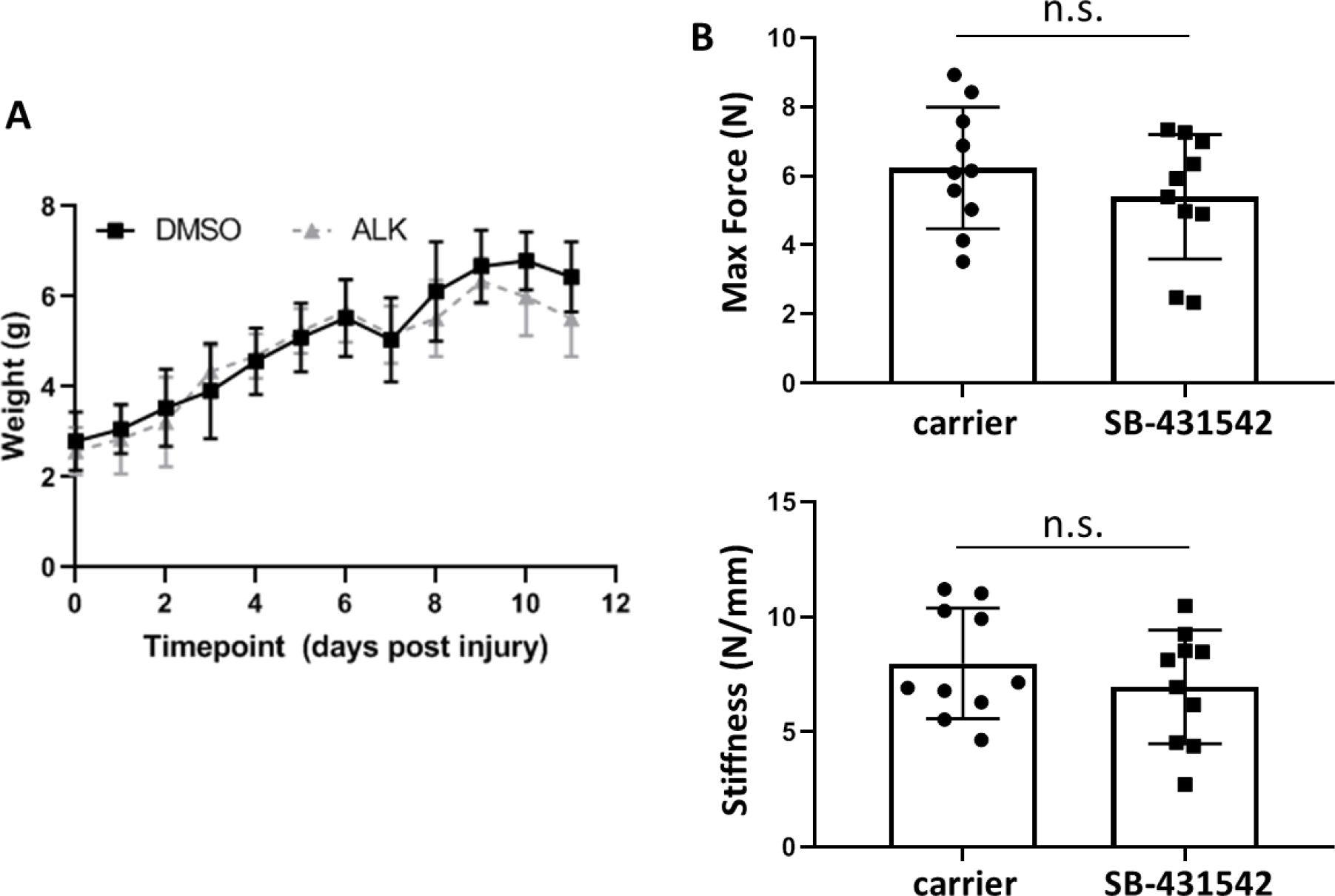
Postnatal growth is not affected by SB-431542 treatment. (**A**) Weight and (**B**) tendon mechanical properties are comparable between carrier-treated and SB-431542-treated mice. n.s. indicates p>0.1.

**Figure S2:**
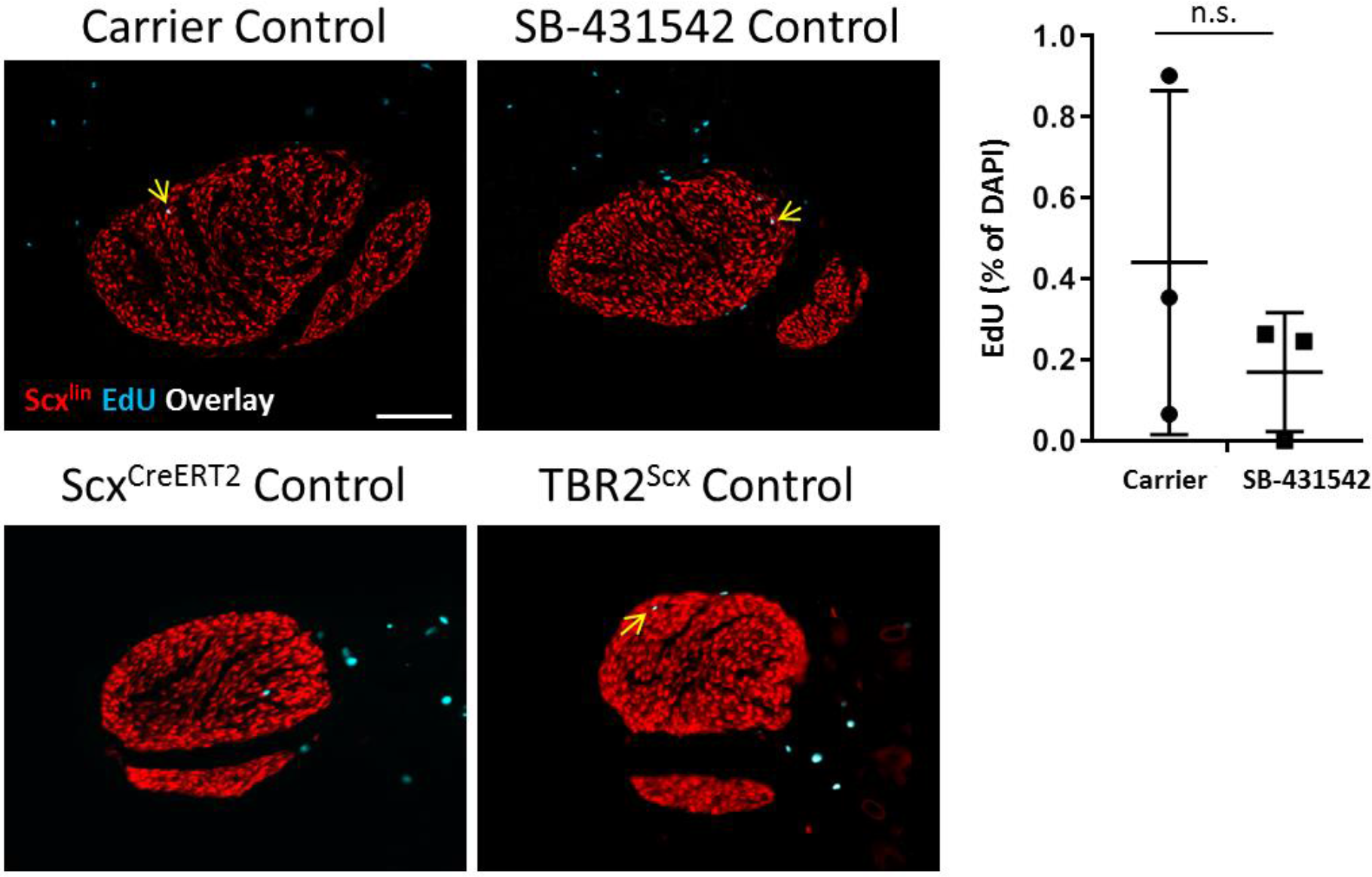
Proliferation in control, uninjured tendons is not affected by SB-431542 treatment or *TβR2*^*Scx*^ deletion. Transverse section images through control tendons stained with EdU show no differences between control tendons with SB-431542 treatment or *TβR2*^*Scx*^. Arrows indicate EdU+, *Scx*^*lin*^ cells. Scalebar: 100 μm.

**Figures S3:**
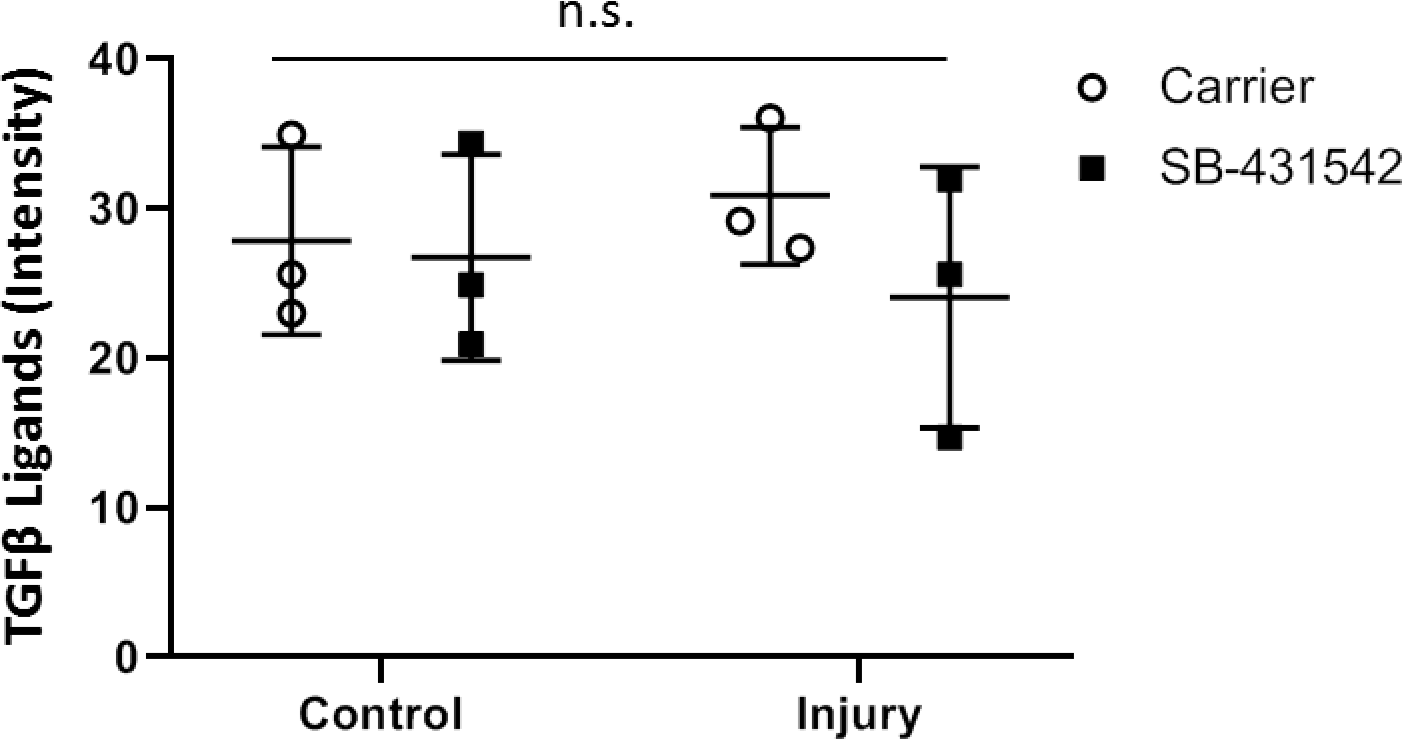
TGFβ ligand within the gap space is not affected by SB-431542 treatment at d3. Quantification of TGFβ ligand immunostaining show no differences in intensity in the gap space with SB-431542 treatment (n=3). n.s. indicates p>0.1. Scalebar: 100 μm.

**Figure S4:**
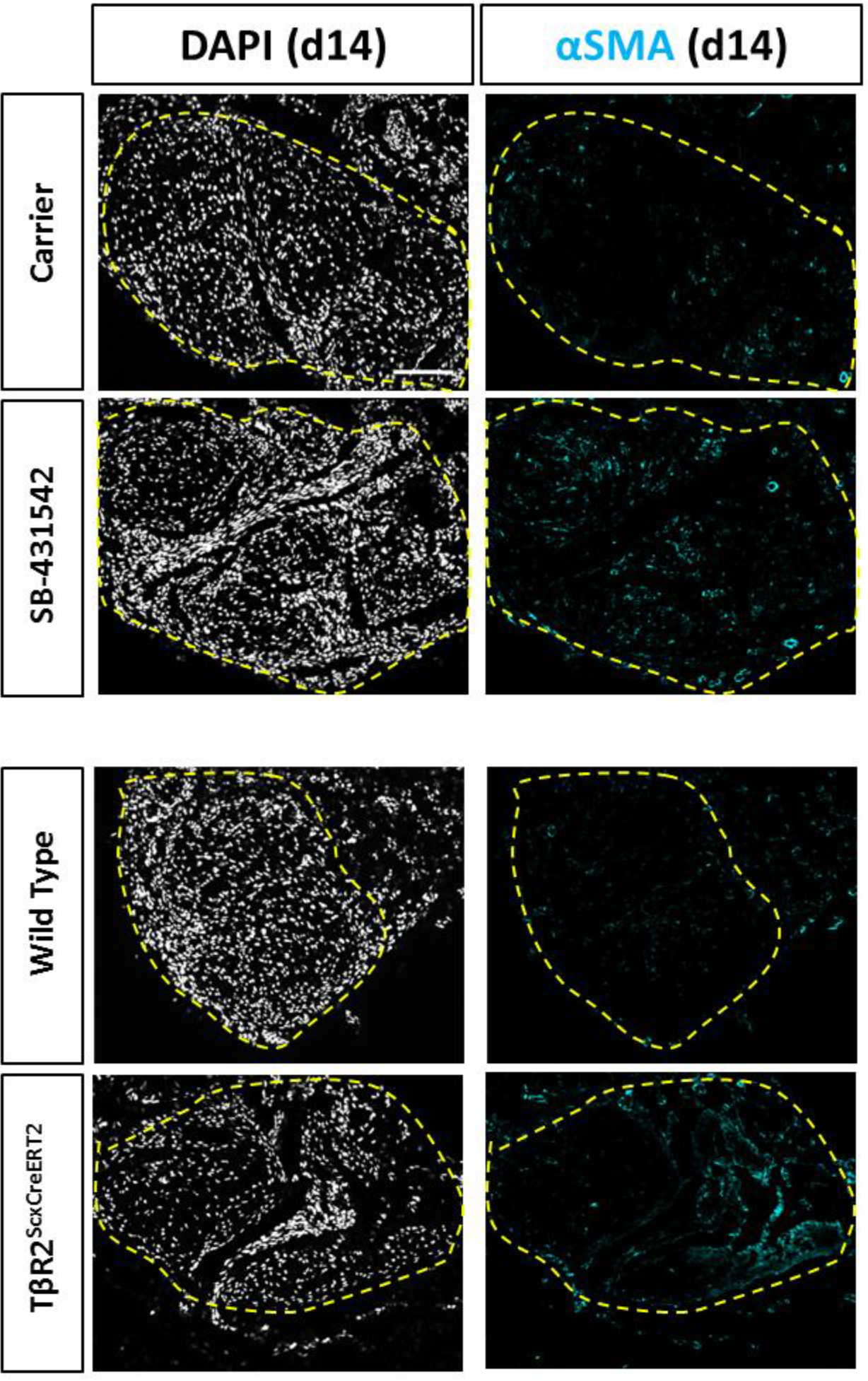
αSMA+ cells are minimally detected by d14 post-injury. Transverse sections stained for αSMA show little staining for all samples. Dashed yellow outlines indicate neo-tendon region. Scalebar: 100 μm.

**Figure S5:**
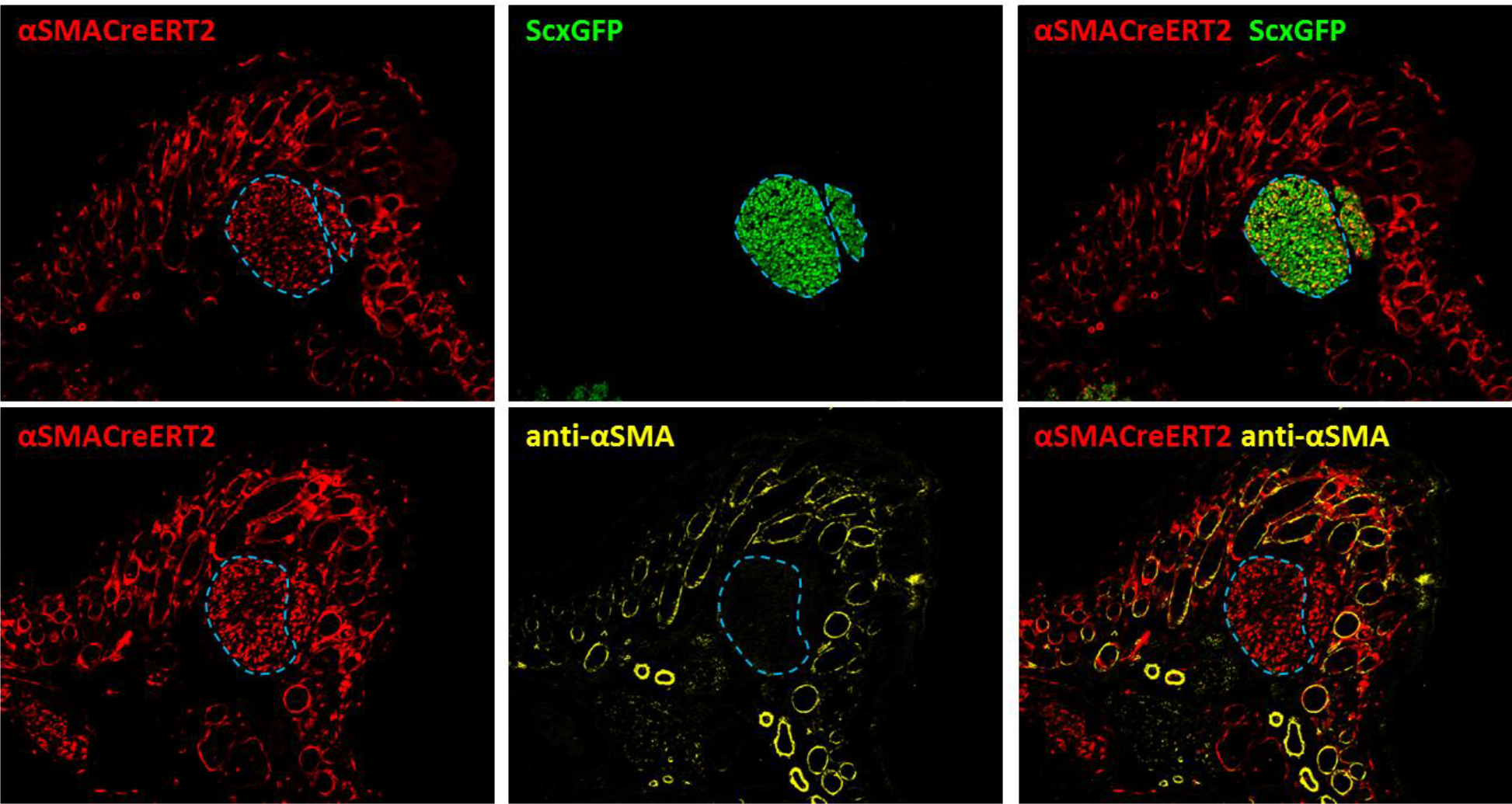
Lineage tracing with αSMACreERT2 show unexpected labeling in neonatal tenocytes. Tamoxifen delivered at P2, P3 with P5 harvest show extensive aSMAlin labeling in ScxGFP+ tenocytes (blue dashed outlines). Immunostaining for αSMA showed that tenocytes do not express αSMA. Overlays show extensive αSMAlin labeling and immunostaining in hair follicles (orange arrows) while blood vessels are inconsistently labeled (green arrows).

